# Critically endangered franciscana dolphins in an estuarine area: fine-scale habitat use and distribution from acoustic monitoring in Babitonga Bay, southern Brazil

**DOI:** 10.1101/2022.10.17.512605

**Authors:** Renan L. Paitach, Guilherme A. Bortolotto, Mats Amundin, Marta J. Cremer

## Abstract

Franciscana dolphins in Babitonga Bay represent the only population of that critically endangered species which is confined to an estuary. Surrounded by large cities and harbors, that environment presents intense human activities and potential impacts that may threaten the dolphins. Understanding their habitat use and distribution can inform mitigation of such impacts. Here we used acoustic data from sixty fixed passive acoustic monitoring stations, implemented between June and December 2018. The relationship between the occurrence of franciscanas and environmental variables was investigated with generalized additive mixed models. The selected model presented 51% of explained deviance and included “time of day”, “intensity of presence of Guiana dolphins”, “maximum slope”, and “bottom sediment”, among other less statistically significant variables. A daily distribution pattern was identified, with franciscanas remaining in the areas of greatest occurrence especially in the morning and seemed to prefer sandy bottom and flatter areas. Areas intensively used by Guyana dolphins were avoided. Additionally, we mapped their distribution using “Empirical Bayesian Kriging” to identify the main areas of occurrence and for foraging. Franciscanas are consistently predominant in the innermost region of the estuary, without expressive use of the entrance channel, but with a wider range in winter than in the spring. The area around the islands, between the north and south banks, represents an important foraging area, a behavior more frequent during dawn and night. This study provides important insights into critical habitat and behavioral patterns of franciscanas, especially this critically endangered population.

## 1. INTRODUCTION

Information on habitat use and distribution of wild animal populations can guide management of conflicting human activities, allowing promotion of conservation strategies (Hastie et al. 2003, Cañadas et al. 2005). Because designating the entire distribution ranges of highly mobile species, such as marine mammals, as protected areas can be practically impossible, identifying priority areas which are essential to their survival, such as those used for foraging and breeding, is of great importance (Hoyt 2012).

The franciscana dolphin (*Pontoporia blainvillei*) is endemic to the Southwestern Atlantic Ocean, occurring from Espírito Santo, in Brazil (18°25′ S), to the Argentinian Patagonia (42°35′ S; Crespo 2009). With high risk of extinction mainly due to high accidental mortality from entanglement in fishing nets (Pinheiro & Cremer 2003), the species is listed as “Vulnerable” globally by the IUCN (Zerbini et al. 2017), and “critically endangered” in Brazil (MMA 2014). Their habitat use is poorly known, with information limited to how that relates to bathymetry, with coarse spatial resolution and temporal dynamics have not yet been considered (e.g., Danilewicz et al. 2009, Amaral et al. 2018, Sucunza et al. 2019). The species is mainly found in coastal habitats on the continental shelf, between the surf zone and the 50 m depths, predominantly up to 30 m deep (Danilewicz et al. 2009), but some individuals are occasionally seen visiting bays and river deltas (Bordino et al. 1999, Di Beneditto et al. 2001, Azevedo et al. 2002, Failla et al. 2004, Santos et al. 2009, Zappes et al. 2018).

The only known distinct franciscana population residing exclusively in an estuarine habitat is found in Babitonga Bay, southern Brazil (Cremer & Simões-Lopes 2008, Cremer et al. 2018). With about only 50 individuals, there is evidence of a high degree of isolation, corroborated by satellite telemetry data, photo-identification and genetic analyses (Dias et al. 2013, Sartori et al. 2017, Cremer et al. 2018, Wells et al. 2021). This population is considered by experts as a demographically independent management unit for conservation purposes (sensu Moritz 1994). In addition to accidental catches in gillnets (Pinheiro & Cremer 2003), habitat degradation by chemical pollution (Alonso et al. 2012), and the construction and expansion of ports, which includes underwater blasting work and dredging, severely compromise the health of the bay’s ecosystem and, consequently, the survival of this dolphin population (Cremer et al. 2018, Paitach et al. 2019).

Visual surveys have indicated a heterogeneous distribution of franciscanas in Babitonga Bay, which are restricted to the innermost regions of the estuary and concentrated around the islands in its central portion (Cremer & Simões-Lopes 2008, Paitach et al. 2017, Cremer et al. 2018). Factors already recognized to influence their habitat use include variations in tidal cycles, which probably reflects prey availability fluctuation, and the presence of sympatric Guiana dolphins (*Sotalia guianensis*) (Paitach et al. 2017, Cremer et al. 2018). Joint conservation strategies for sympatric species, ecologically similar and that share limited resources, can benefit from the understanding of how such species affect or influence each other (Bearzi 2005).

Understanding the ecological requirements of small cryptic cetaceans is a major challenge, standard visual surveys are not always an option. Franciscanas are one of the smallest dolphin species, form small groups, rarely display aerial behaviors, and only expose a small part of the body when breathing at the surface for a very few seconds (Wells et al. 2013, Cremer et al. 2018; Actis et al. 2018). Furthermore, visual observations, whether from vessels or aircrafts, are restricted to daylight periods and require very good weather conditions. Like most cetaceans they produce sounds when diving, which allows acoustic sampling (Tyack & Clark 2000).

Passive acoustic monitoring (PAM) allows the autonomous logging of the underwater sounds generated by cetaceans and can be an efficient alternative to visual surveys for detecting their presence (Paitach et al., 2021). PAM can be used to investigate various ecological and behavioral aspects of cetaceans (Batista & Gaunt 1997), can record in bad weathers and during the night, and have low associated costs (Mellinger et al. 2007, Van Parijs et al. 2009). Cetacean echolocation click trains detected in PAM stations distributed in an area of interest, for example, illustrate how PAM can be used to identify potential foraging areas and periods (e.g., Pirotta et al. 2014, Tubbs et al. 2020, Paitach et al. 2021). PAM have been widely used worldwide for studies on cetacean distribution, migrations, behavior, habitat use, and identification of impacts and threats (e.g., Verfuß et al. 2007, Mellinger et al. 2007, Van Parijs et al. 2009, Jaramillo-Legorreta et al. 2016, Carlén et al. 2018). For franciscanas, it was only used to describe their acoustic repertoire and sound production characteristics (Tellechea et al. 2017, Barcellos & Santos 2021, Paitach et al. 2021).

We used an array of PAM devices for sampling franciscanas sounds during winter and spring in Babitonga Bay. Our objectives were to identify the main environmental variables related to how that population use the habitat in the Bay, including the presence of Guiana dolphins, and to produce maps of franciscanas’ distribution and foraging areas that could inform conservation strategies and management of human activities. We hypothesized that franciscanas vary their distribution between seasons and along the day, and that the patterns are linked to environmental features and niche partitioning with Guiana dolphins.

## 2. METHODS

### 2.1. Study area and sampling design

A systematic grid was designed for deploying sixty PAM stations in Babitonga Bay (26°02’-26°28 ’S – 48°28’-48°50’ W), Santa Catarina State, southern Brazil (Fig. 1). The study area is approximately 160 km^2^ wide, with 6 meters average water depth, and some extremely shallow areas, which become exposed at low tide (Vieira et al. 2008). The waters in the bay are supplied from several rivers, but its physical-chemical characteristics are spatially homogenous (IBAMA 1998). It has a semi-diurnal regime of micro tides, meaning two well-defined daily cycles of floods and ebbs during spring tides, reaching a maximum amplitude of less than 2 m (Vieira et al. 2008). Since the grounding of the narrow southern channel for construction of an access road to the São Francisco do Sul Island in 1937 (thick black segment in Fig. 1), the only connection to the open ocean is through a 28 m deep channel to the north.

**Figure 1.**
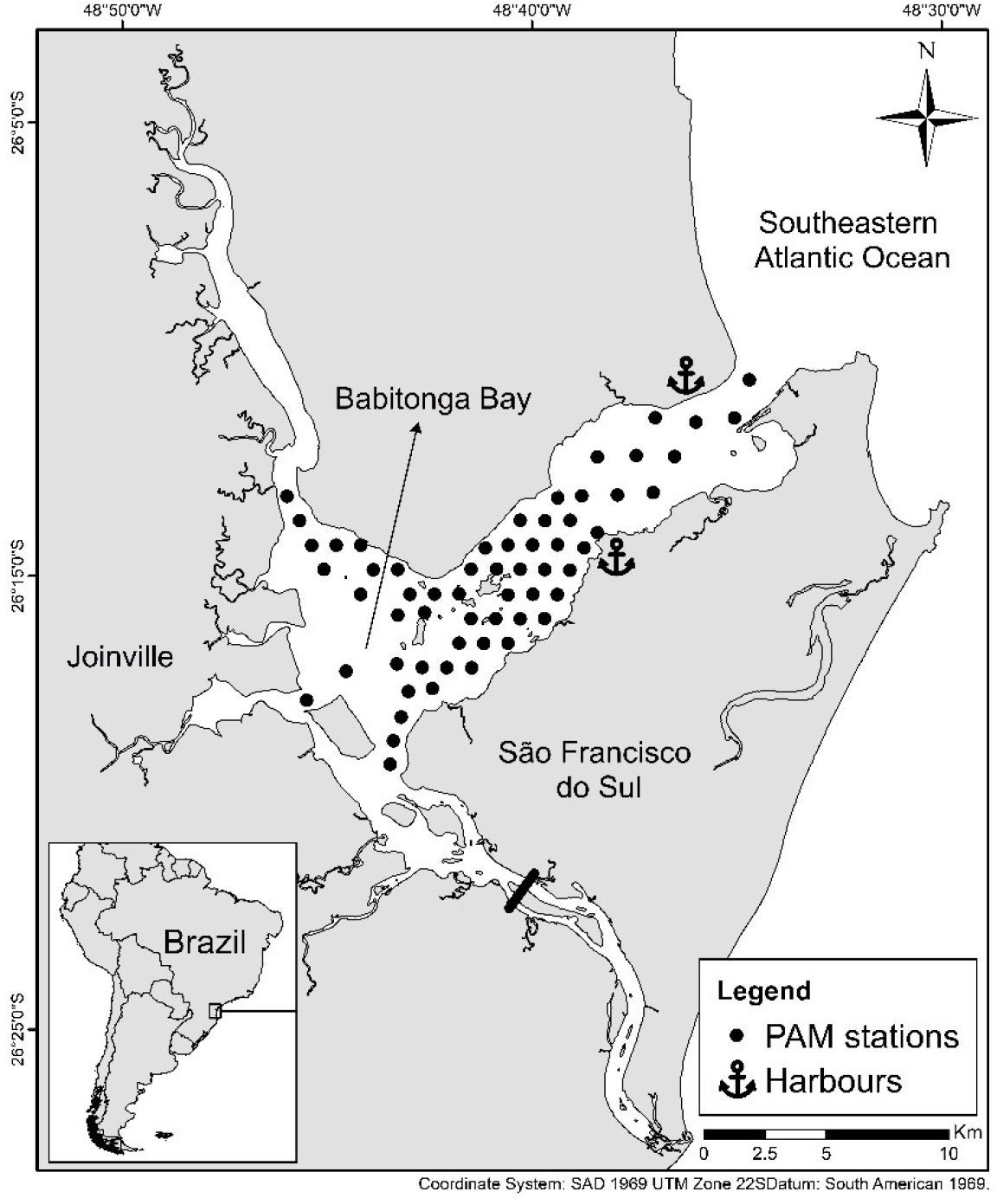
Distribution of sixty passive acoustic monitoring (PAM) stations deployed in Babitonga Bay, southern Brazil, for recording franciscana dolphins. A think black segment indicates the location where the historical south channel was grounded in 1937 for the construction of an access road to the São Francisco do Sul Island.

### 2.2. Sampling methods

Acoustic monitoring was done using C-PODs (Chelonia Limited©, UK), i.e., autonomous acoustic loggers designed to log trains of tone-like pulses between 20 and 160 kHz. Such devices are ideal to record narrow-band high frequency (NBHF; i.e. peak frequency at 130 kHz and no essential energy below 100 kHz) sonar click trains of franciscanas, but also the broadband clicks of Guiana dolphins (Paitach et al. 2021). C-PODs have an omni-directional hydrophone (i.e., records in all directions) and with a source level of 190 dB and a detection threshold of 120 dB, the theoretical detection range of a franciscana should be around 400 m on-axis (Nick Tregenza, Chelonia Ltd., personal communication, 2017). They were fitted into custom-made cages, designed to protect them from net entanglement (Paitach et al. 2021). Cages had none or negligible interference in the acoustic recordings. PAM stations sites were determined at semi-random within the survey area, constrained to locations with a minimum average depth of 4 m, and were at least 1,600 m apart in the access channel and 800 m in other areas (Fig. 1) to proportionally address the expected distribution of franciscanas in those areas (c.f. Cremer et al. 2018); i.e., the more concentrated the stations, the more dolphins expected.

Acoustic samples were collected from June 26 to December 24, 2018, with a varying number of days monitored at each station. In total, 35 C-PODs were used, with a maximum of 20 C-PODs operating simultaneously. A subset of C-PODs (usually 20) was replaced every approximately 30 days by others with fresh batteries and SD cards, always using new PAM station. To minimize systematic bias from possible differences in C-PODs detection potential due to temporal degradation (Dähne et al. 2013), devices were randomly placed at each deployment, as recommended by Carlén et al. (2018). The design aimed at sampling each position for 30 days in the winter and 30 days in the spring, on average. Ten C-POD subsets were defined with three station positions each, considering the closest possible positions for each group, and each of these positions was sampled at each exchange, ensuring that the distribution of the monitored points remained as homogeneous as possible in the area over the study period.

### 2.3. Data analysis

Franciscana sonar click trains were identified using KERNO click train classifier in CPOD.exe (Chelonia Limited, UK). That software identifies NBHF-type sounds with higher robustness and lower levels of false positives than classifiers based on individual clicks (Dähne et al. 2013, Roberts & Read 2014). Only click trains classified by KERNO as having a “high” or “moderate” probability of being generated by franciscanas were analyzed. The “Detection Positive Minutes” per hour (DPM/h; number of minutes with at least one franciscana click train within an hour) were extracted and was used here as a proxy for the intensity of franciscanas presence.

The C-POD has a limited of 4,095 logged pulses per minute to avoid data overload and, consequently, saturation of the memory card and battery consumption. After that limit, the logging is interrupted and only resumed in the following minute. Ambient noise, such as the sound of rain, moving bottom sand, or produced by living organisms such as shrimp and fish, all of which may generate pulsed sounds that can be logged by the C-PODs. Excessive noise data were evaluated with the ‘Detections and Environment’ tool in CPOD.exe and disregarded to ensure sampling homogeneity.

### 2.4. Habitat use

For modelling of franciscanas’ habitat use on a fine scale, the intensity of their presence around PAM stations, expressed as DPM/h, was modelled as a function of environmental variables. Generalized Additive Models (GAMs; Hastie & Tibishirani 1990) were applied using R software v.4.0.3 (R Core Team 2020) to accommodate the possibly complex relationships between franciscanas occurrence and variables. Because the data set to be modelled was large (n = 64,745), models were fitted using function “bam” (mgcv R package; Wood 2017) which allows relatively fast model fitting. Preliminary inspection was conducted to ensure that the data contained information useful for inference on habitat use. Maps and graphics illustrating sampling distribution balance over time confirmed the adequacy of the data (Supp. material). The negative binomial distribution showed the best fit and was adopted for modelling.

Environmental variables (Table 1) were obtained for each PAM station using ArcGIS Pro 2.3 (https://www.esri.com), with input data from morphosedimentary and topographic databases (provided by Vieira et al. 2008). Tidal conditions for each monitoring hour were attributed to PAM stations using tide tables published by the Directorate of Hydrography and Navigation of the Brazilian Navy for the port of São Francisco do Sul. The DPM/h of the Guiana dolphin was also included as a variable. The classification procedure for this species was like that for franciscanas, with virtually zero risk of miss-specification (see Paitach et al. 2021).

**Table 1.** Variables used for modelling habitat use of franciscana dolphins in Babitonga Bay.

| Variables | Range of values | Explanation and Categories |
| --- | --- | --- |
| UTMX | 723237 – 741747 | Longitude in UTM |
| UTMY | 7086720 – 7101381 | Latitude in UTM |
| Hour.of.day | cyclic | 24-hour circadian cycle |
| Month | 6 – 12 | Months of the year, from June (6) to December (12) |
| Tide.state | categorical | Tidal cycles: flood, high, ebb and low |
| Tide.type | categorical | Type of tidal amplitude as a function of the Sun-Moon gravitational conjunction: syzygy = full and new moons; quadrature = first quarter and third quarter moons |
| Sg.DPM | 0 – 60 | Detection Positive Minutes of *Sotalia guianensis* per hour |
| Season | categorical | Austral seasons: winter = from June 20 to September 21; spring = from September 22 to December 20 |
| Carbonate | categorical | Percentage of inorganic salts in the sediment within a radius of 400m: 0-10%; 10-20%; 20-30%; 30-40% |
| Organic.matter | categorical | Percentage of organic matter in the sediment within a radius of 400m: 0-2%; 2-4%; 4-6%; 6-8%; 8-10% |
| Sediment | categorical | Predominant texture of the bottom sediment within a radius of 400m: sand; sand with mud; mud with sand; mud |
| Deep.max | 2 – 22.3 | Maximum depth in meters within a radius of 400m |
| Deep.min | 0.1 – 6.9 | Minimum depth in meters within a radius of 400m |
| Deep.mean | 1.8 – 10.7 | Average depth in meters within a radius of 400m |
| Deep.range | 1.5 – 18.1 | Range between minimum and maximum depth within a radius of 400m |
| Slop.mean | 0.179 – 3.364 | Average slope in degrees within a radius of 400m |
| Slope.max | 1.519 – 51.388 | Maximum slope in degrees within a radius of 400m |
| Aspect | 59.096 – 258.092 | Average direction of the slope in degrees from north within a radius of 400m |
| TCI | 0.0001 – 0.6613 | Topographic complexity index calculated by multiplying scaled values for slope and aspect (Bouchet et al. 2015) averaged within a radius of 400m |
| Margin.distance | 146.5 – 1952.5 | Distance in meters from the nearest margin. |
| Margin.feature | categorical | Feature of the nearest margin: continent or island |

Linear correlation and concurvity, a measure of non-linear relation between smooth terms within a GAM, were verified for a preliminary model which including all available variables. All measures of depth (Table 1) were linearly correlated to each other, to slope measures and to geographic location, UTMX and UTMY. Aspect and TCI were linearly correlated to each other. Correlated variables were not included in the same model.

Preliminary models indicated that residual autocorrelation could be a problem. Correlation structures presented a cyclic pattern with an apparent peak every 24 units apart. To account for that, a 2-D smoother (Wood 2017) for easting (i.e., “UTMX”) and northing (i.e., “UTMY”) combined, with a different tensor for each hour of the day, was added to all models. That approach allowed the spatial heterogeneity in the data to be explicitly modelled as a function of time and space. Also, a first-order autoregressive error structure function (AR1) was added in the models. For each model, the AR1 correlation parameter *ρ* was calculated by fitting models without correlation structure and measuring the first lag in the autocorrelation function (“acf”, R function). In the present modelling framework, the AR correlation structure corresponded to a GEE (Generalized Estimating Equations; Ziegler, 2011) approximation which, in practice, increased the uncertainty in the estimated smoothers. That means that p-values for smooth terms became larger when compared to corresponding models without AR1 structure. Since the data set was formed by time series, with observations representing repeated measurements for each location, a smooth term for each sampled PAM station as a random variable was used in all models.

Smooth functions were used to model the relationship between continuous variables and the response value. Except for the 2-D smoother for easting and northing combined with a tensor for each hour of the day and a cyclic spline for hour of day (“Hourday”), thin plate regression splines were used (R package “mgcv”; Wood 2017). The dimension basis (i.e., parameter *k* on smooth functions, mgcv R package) was set to a maximum of seven for all tested smoother of variables, to both avoid overfitting and prevent smooth functions impossible to interpret biologically. For variables “Aspect” and “Maximum Slope”, that parameter was further decreased to five, because preliminary modelling showed fitted smoothers of hard biological interpretation, i.e., with several peaks.

Model variables were selected in a forward step approach, based on minimum Akaike Information Criterion (AIC; Akaike 1974): the initial model presented a 2-D smooth function for UTMX and UTMY with a different tensor for each hour of the day, a smooth function for “Point” as a random variable, and a cyclic smooth term for “Hour of day”. In the first round of variable selection, models with only one additional variable were fitted, and the one presenting the smallest AIC score was considered as the initial model in the following step. In each step, only one additional variable was separately added to the model selected in the previous step. Those steps were repeated until the AIC could not be improved by the addition variables, and so the resulting model was retained as the most efficient to describe the variation in the presence of franciscanas.

### 2.5. Distribution

The distribution of franciscanas and of their foraging activity were investigated through interpolation of spatial data (i.e., “kriging”) using software ArcGIS Pro 2.3 (Geostatistical Analyst; Geostatistical Wizard; https://www.esri.com). Kriging is a geostatistical interpolation method that assumes that the distance or direction between the points in the sample reflects a spatial correlation that can be used to explain the variation in the surface (Oliver & Webster, 1990). Without imposing a priori environmental variables, the spatial autocorrelation of a specified number of points is modeled in semi-variograms which are used to estimate density at each location (Oliver & Webster 1990).

More specifically, Empirical Bayesian Kriging (EBK) was used. While other kriging methods require several projection parameters to be manually adjusted, EBK automatically calculates these parameters at each predicted location using a subset process and data simulations. The method also differs from other kriging methods by taking the standard error introduced by the estimate of the underlying semi-variogram into account, propagating that uncertainty when generating predictions in locations not surveyed (Oliver & Webster 1990, Krivoruchko 2012). Semi-variogram parameters were estimated using restricted maximum likelihood (REML), which is indicated for small data sets to avoid overestimating densities at restricted areas (Krivoruchko 2012).

Two variables were separately used to generate distribution maps: 1) Detection Positive Hours (DPH) was used to identify the main areas of franciscana occurrence; and 2) adjusted Feeding Buzz Ratio (FBR) was used to identify foraging areas. The DPH was obtained using the KERNO classifier and a similar selection criteria as the DPM used in the analysis of habitat use, however with hours as period of interest (i.e., coarser temporal resolution). All click trains recorded throughout the study were exported and classified as “feeding buzzes”, based on an Inter-Click Interval (ICI) of less than 10 ms (Carlström 2005, Paitach et al. 2021). FBR values were then calculated as the ratio between number of buzzes and number of non-buzz click trains (with ICI > 10ms). A weighted metric of the importance of the foraging areas was obtained by adjusting FBRs by the intensity of franciscanas occurrence (i.e., multiplying the FBR by the DPH).

Seasonal (winter and spring) and diel (dawn = 00:00-05:59, morning = 06:00-11:59, afternoon = 12:00-17:59, and night = 18:00-23:59) maps were produced. The midday and midnight cut-off limits were chosen to allow a some understanding for distribution patterns within the light and dark periods. Those periods can be more easily used for illustrating management strategies related to the time of the day. Average values of DPH and adjusted FBR were calculated separately for each day (for season maps), and for each period of the day (for diel period maps), and then averages for all sampled days were calculated for each PAM station. Days with less than 24 hours of data collected or periods of the day with less than 6 hours collected were not considered in this analysis. Since the FBR values are adjusted, biological interpretation can be difficult. Therefore, maps for FBR values were grouped into classes of importance. Outliers were removed and the resulting scale of values was divided into four equally sized classes. The lowest class was disregarded (low importance), and the others were ‘moderate’, ‘high’, and ‘very high’ importance for foraging.

## 3. RESULTS

Out of the 60 monitoring stations planned throughout the study area, only 6 were not sampled in winter, and 11 in spring, due to loss of equipment. PAM stations were monitored for an average of 28 days (minimum of 3, and maximum of 57 days) in winter and 24 (minimum of 2 and maximum of 91) in spring. A total of 66,350 hours of acoustic recordings were collected in 182 days, both seasons considered. After data filtering (i.e., removing data with excess noise) 64,745 hours were analyzed, 7,432 (11.5%) of which with franciscana recordings.

### 3.1. Habitat use

The final habitat use model had 51% of explained deviance and fitted the data well, except for high values of the response variable (Supp. material). Despite the assumption of residual constant variance not being fully met, the negative binomial distribution (θ = 0.092) showed the best fit to the residuals. Residual autocorrelation was greatly reduced by the inclusion of an autoregressive function in the model, yet still mildly present (Supp. material). For that reason, the inclusion of variables in the final model must be interpreted carefully, especially for variables with lower significance (i.e., large p-values). Coefficients for factor variables and smooth functions included in the final model can be checked in the Supplementary material.

The forward step variable selection resulted in the inclusion of smooth functions for intensity of presence of the Guiana dolphins (“SG.DPM”) and maximum slope (“Slope.max”), in addition to the compulsory smother in the initial model (i.e., “Point” as a random variable; a 2D smoother for “UTMX” and “UTMY”, with a tensor for each hour of the day; and a cyclic smoother for “Hour.day”) (Fig. 2). There was a clear cyclic pattern in the occurrence of franciscanas across the study area, indicating that in the areas where their occurrence was more intense, they were more likely to occur during the early hours of the day. Areas with very high values of intensity of presence of Guiana dolphins were avoided by the franciscanas, but to a lesser extent, they seemed to be tolerated. Franciscanas seem to avoid steeper areas within the range of slopes in Babitonga Bay.

**Figure 2.**
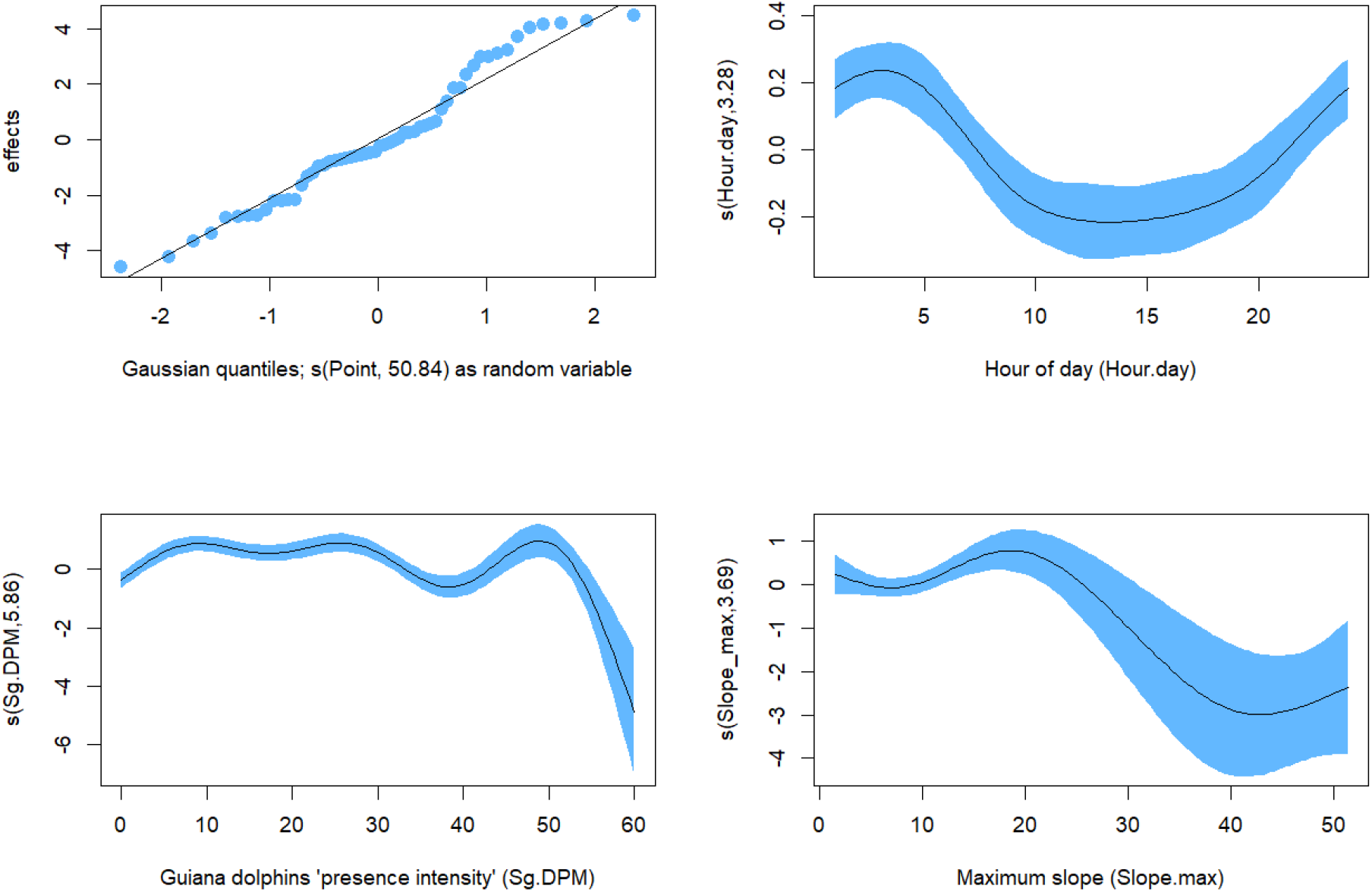
Smooth functions for variables included in the final model for habitat use of franciscana dolphins in Babitonga Bay. Degrees of freedom are shown inside parentheses.

The final model also included factor variables “Month”, “Sediment”, “Tide.type” and “Tide.state”. Because of multiple factor variables, partial effects for each combination of factor levels would require several plots. Boxplots of values adjusted for the intensity of the presence of franciscana (Pb.DPM) for each selected factor variable are shown individually (Fig. 3). The presence of franciscanas seems to vary slightly over the months of study, but a clear seasonal pattern was not observed. The presence of franciscanas in Babitonga was associated with the granulometry of the bottom sediments, with a greater presence over sandy bottoms and less presence over mud bottoms. Despite contributing to improving the model AIC, it is not clear how tide variables were related to the variations of presence of franciscanas, since the levels were not precisely estimated, as indicate by large p-values (Supp. material).

**Figure 3:**
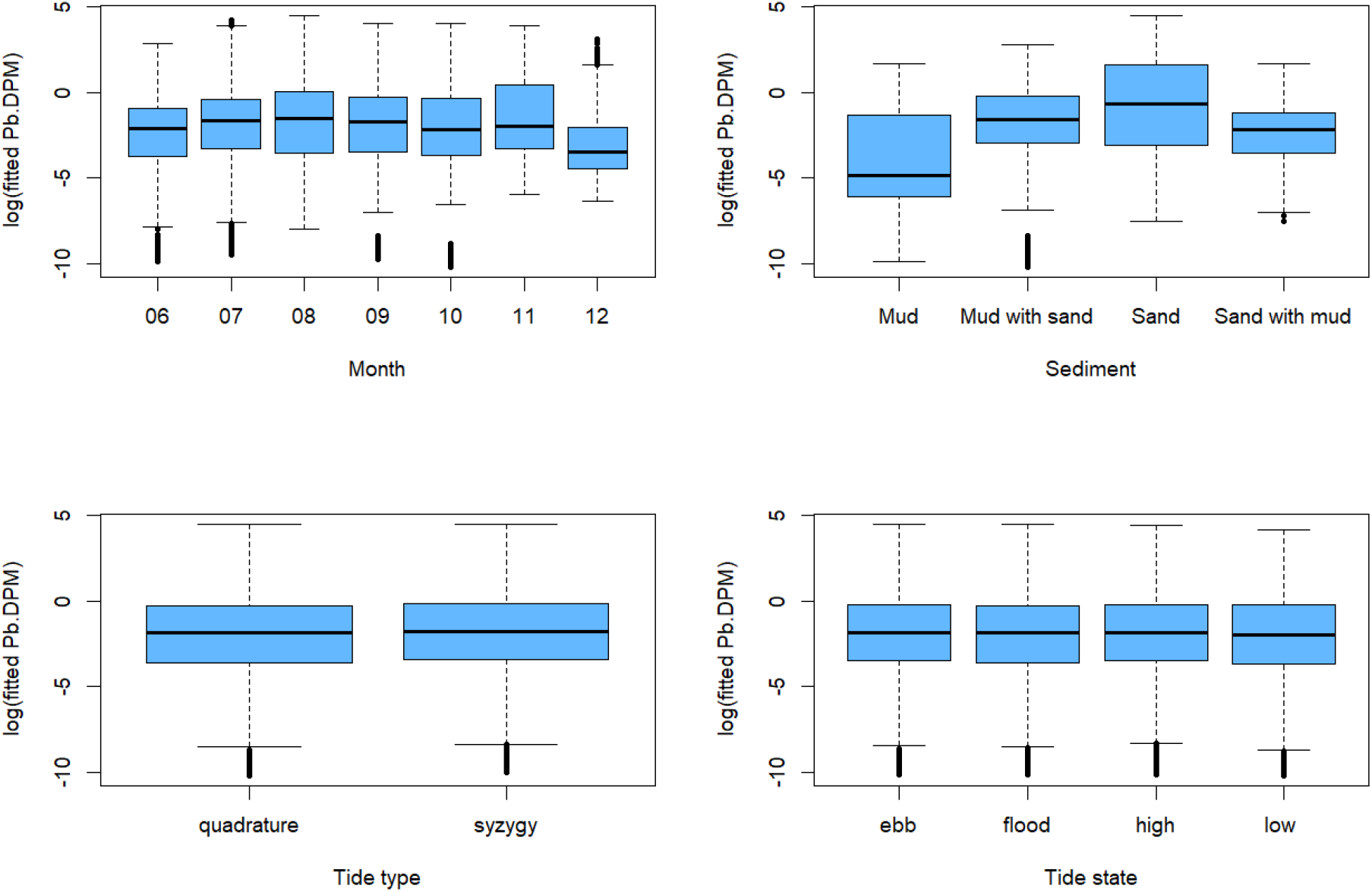
Boxplots for fitted values in the final model for different levels of the factor variables included in the final model for habitat use of franciscana dolphins in Babitonga Bay.

### 3.2. Distribution

Predictive maps of occurrence and foraging areas were generated for each season (Fig. 4). The distribution of franciscanas was predominant in the innermost region of the estuary, without a marked use of the open sea access channel. In the winter their distribution expanded, extending to the mouth of the Palmital River (northwest axis), the entrance to Saguaçú Lagoon (west margin), and the Linguado channel (south axis), and further along the northeast margin of the bay. In the spring the distribution was predominantly in the central region of the bay, between the north and south margins. The area with the highest density in winter was located slightly towards the west than in spring, which remained closer to the north-central margin. The area between the north margin and the islands represents important franciscana foraging areas, both in winter and spring, but in winter the area between the islands and the south margin were also important for foraging. In winter, the northeastern margin, and the area close to the mouth of the Palmital River (northwest axis) also appear to be areas used for foraging, which were not seen in the spring.

**Figure 4:**
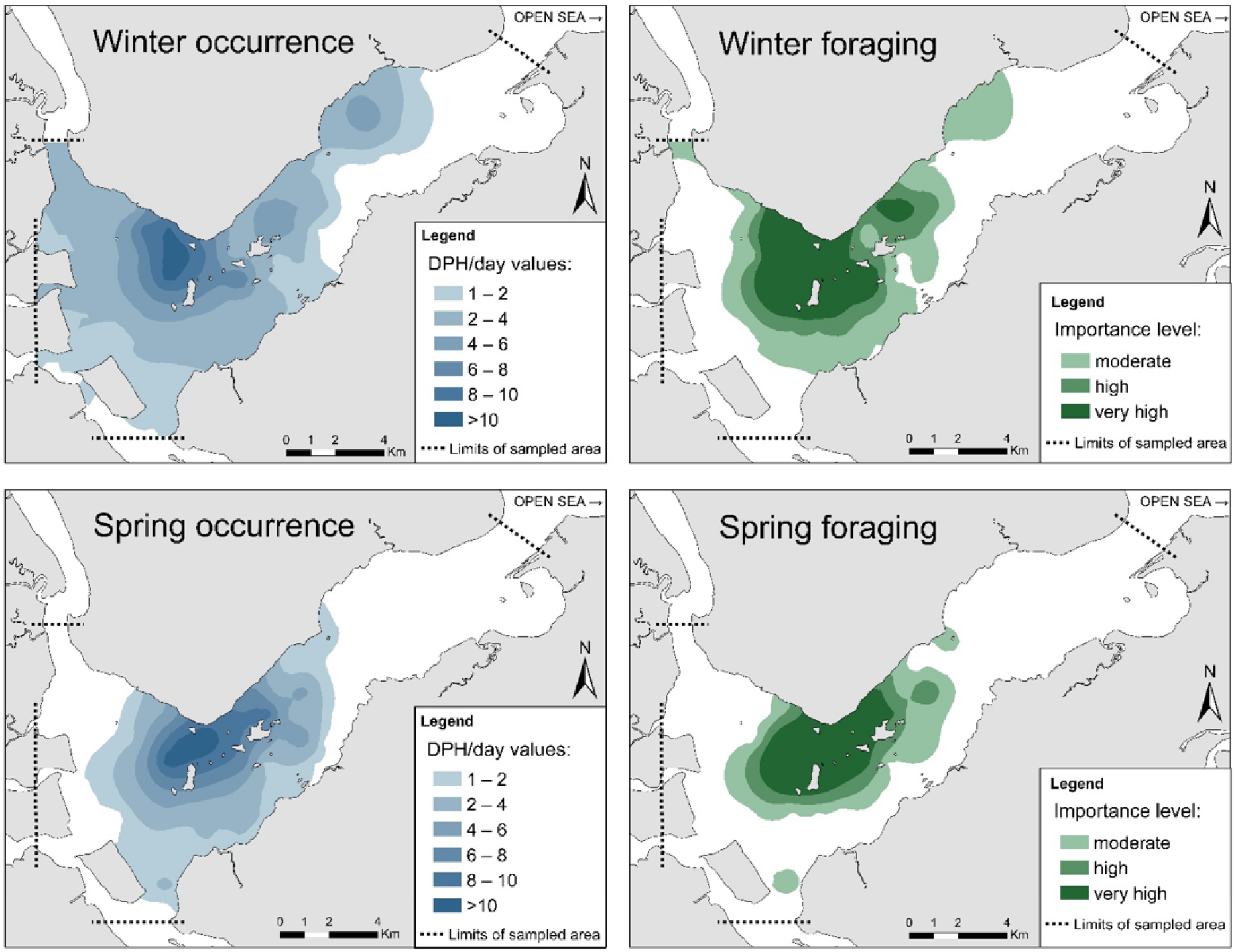
Winter and spring occurrence and foraging areas of franciscana dolphins in Babitonga Bay. (DPH/day = detection positive hours per day). Foraging importance level estimated by multiplying DPH/day by the Feeding Buzz Ratio (see text for details).

Areas of occurrence and important for foraging for franciscanas varied throughout the diel periods in both seasons (winter: Fig. 5; spring: Fig. 6). The central area of the bay, between the islands and the north margin, remained as the core area of franciscanas throughout the day, in both seasons, while areas with less intensity of use varied throughout the day in each season.

**Figure 5:**
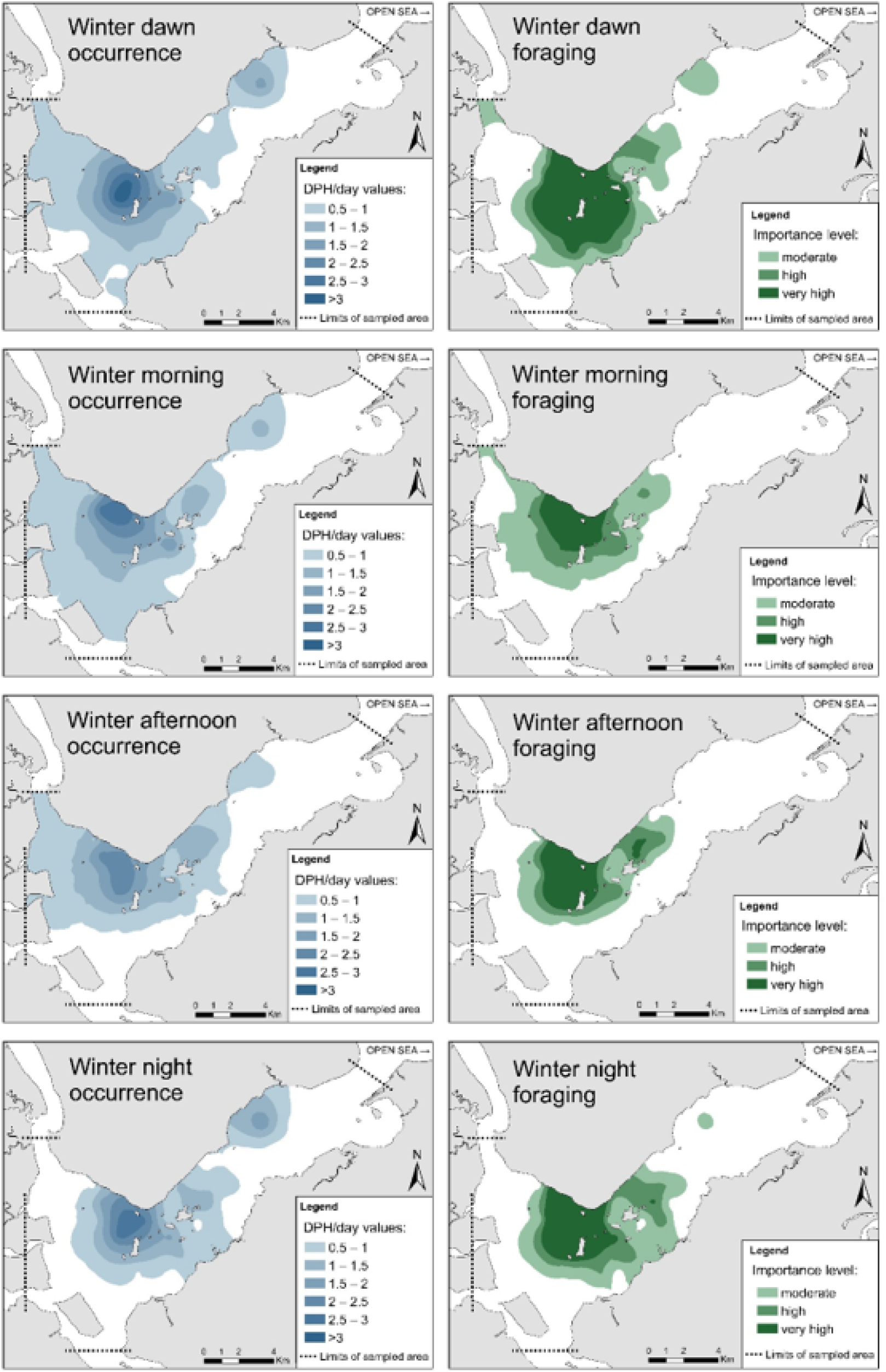
Winter occurrence and foraging areas of franciscana dolphins in Babitonga Bay for each period of the day (dawn = 00:00-05:59, morning = 06:00-11:59, afternoon = 12:00-17:59, and night = 18:00-23:59). (DPH/day = detection positive hours per day).

**Figure 6:**
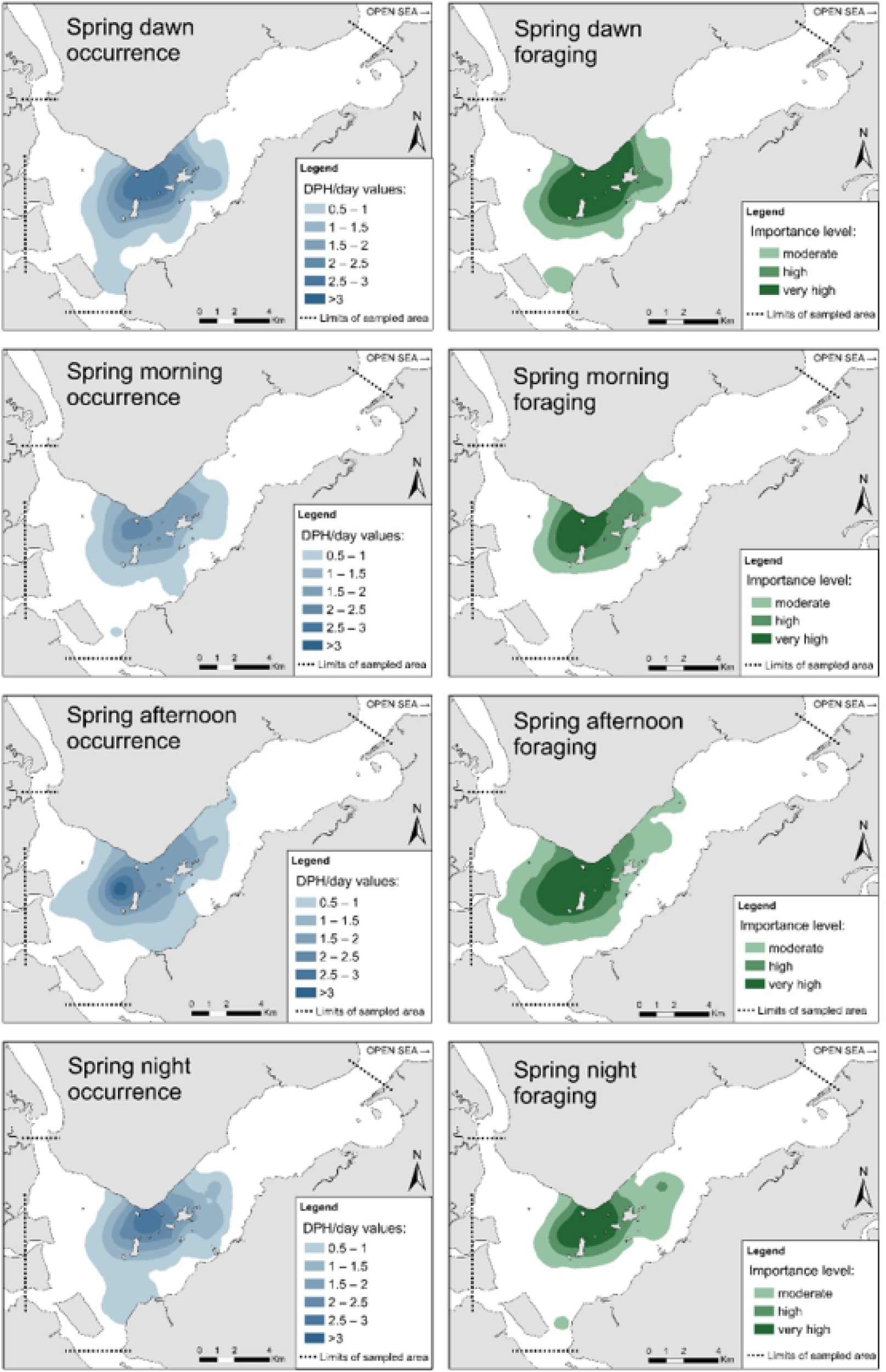
Spring occurrence and foraging areas of franciscana dolphins in Babitonga Bay for each period of the day (dawn = 00:00-05:59, morning = 06:00-11:59, afternoon = 12:00-17:59, and night = 18:00-23:59). (DPH/day = detection positive hours per day).

The distribution of important areas for foraging fluctuated throughout the diel periods in different seasons. In winter, foraging was more concentrated near the core area during the morning and afternoon, and at night it expanded southwards, to a region known as the Laranjeiras channel, which was intensified at dawn (Fig. 5). In the spring the foraging areas were more restricted, with some oscillation in the east-west direction (Fig. 6). During the night they expanded eastwards, occupying the entire surroundings of the islands. During the afternoon there the pattern was towards the opposite direction, with foraging in the innermost portion of the bay, up to its west margin, in an extensive area of shallow water and muddy banks (Fig. 6). In both seasons, the dawn period showed the biggest patches of ‘very high’ importance for foraging, indicating that the feeding behavior is more intense in that period, followed by the night in winter and the afternoon in spring (Fig. 5 and 6).

## 4 DISCUSSION

### 4.1. Passive acoustic monitoring: potential and limitations

The PAM approach and especially the use of C-PODs showed promising signs of a very valuable tool for investigating spatio-temporal patterns of habitat use and distribution of franciscanas. This is the first systematic effort of this nature for the species. The processing of the large data volume obtained (more than 66,000 hours) was facilitated through the C-POD system automated procedure, which saving time and also reduce the potential subjectivity bias of the researcher (Rayment et al. 2009).

A key assumption in the present study is that the heterogeneity observed in the franciscana acoustic detections would reflect the density of these animals in the bay. Failure to meet that could rise from when animals are present but not detected, but comparative distribution studies using visual and acoustic detections indicate that, depending on the species, that may not be an issue (Verfuß et al. 2007). Similarly to harbour porpoises (*Phocoena phocoena*) in the wild, that click almost continuously and with maximum silent intervals of less than 15 seconds (Akamatsu et al. 2005), it is very likely that franciscanas also continuously echolocate in the estuarine waters of Babitonga Bay, which presents a complex topography and very high water turbidity with virtually no visibility (Oliveira et al. 2006, Vieira et al. 2008). Furthermore, and because this is a closed population (Dias et al., 2013; Cremer et al. 2018), numbers of acoustic detections in the study area are not expected to be influenced by emigration/immigration of individuals. Also, since areas with an average depth of less than 4 m, potentially dry at low tides, were under-sampled, it is possible that in periods when lower detection numbers were recorded within the sampled area (i.e., where the water was continuously deeper than 4m), animals might have been in those shallower areas.

We assumed a homogeneous probability of detection of franciscanas by C-PDOs over space and time. It is known, however, that sound propagation may be influenced by spatial and temporal variations in the behavior of the dolphins (Verfuß et al. 2009, Leeney et al. 2011), and by environmental conditions, such as water temperature and salinity (Richardson et al. 1995). There is a trade-off between the range and directionality of the sounds produced by dolphins during traveling and foraging behaviors (Tyack & Clark 2000). Understanding how different behaviors can affect detection probability of franciscanas by PAM can assist the accuracy of future studies. Temperature and salinity affect the speed and absorption of sound in water (Richardson et al. 1995, Ainslie & McColm, 1998), but considering a low variation of these parameters in the study area we claim that this bias is negligible.

Despite the protective cages, entanglement in nets became a problem throughout the study, causing the loss of some equipment units, a problem that was intensified during the spring and forced an early ending of survey after six months of start. We recommend that future studies employ more extensive effort into clearly communicating with fishing communities, so that such incidents can be avoided or that the PAM devices are returned in case of undesired misplacement.

The two seasons sampled in the present study, winter and spring, were strategically to identify priority habitats for the franciscanas. The winter is the season of least availability of food (Cremer 2007), and so the franciscana distribution reflect its most critical places for foraging during a period of food scarcity. The protection of foraging areas is essential for small cetaceans, which are particularly vulnerable to environmental impacts that can reduce prey availability, due to their high food requirements and apex position in the marine food webs (Ross et al. 2011, Wisniewska et al. 2016). In turn, spring represents the main birthing period for the population (Cremer et al. 2013). The protection of important breeding areas is essential for the conservation of small cetaceans, since the stages of young life are particularly vulnerable to species threats (Ross et al. 2011).

### 4.2. Cyclic patterns of habitat use

There was a clear diel cyclic pattern in the occurrence of franciscanas across the study area. In the areas where their occurrence was more intense, they were more likely to occur during the early hours of the day (Fig. 3). That is possibly a reaction to environmental cycles, which modify the abiotic conditions of ecosystems, with biological organisms corresponding (Aschoff 2013). Behavior patterns in response to diel cycles can be diurnal, nocturnal or twilight (Fernandez-Betelu et al. 2019). In coastal environments, tidal cycles can also cause environmental changes that can result in periodic movements of many species, including cetaceans (Gibson 2003). Similarly to what happens in Anegada Bay, Argentina (Bordino et al. 1999), the franciscanas in Babitonga Bay were found to present movement patterns related to the tides, moving towards the mouth of the bay during ebb and in the opposite direction during the flood, following the current flow (Paitach et al. 2017). In the present study, although the tide was selected as an important factor for the habitat use, it was not possible to clearly identify a pattern. In fact, the tidal cycles effects on dolphin habitat use patterns can vary seasonally, and cetaceans appear to be less influenced by tides in open areas than in narrow channels (Pierpoint 2008, Fernandez-Betelu et al. 2019).

Franciscanas seem to avoid steeper areas within the range of bottom slopes in Babitonga Bay. This may be linked to bathymetry also, since depth variables were not included because of their correlation with geographic location (i.e., UTMX, UTMY). Holz (2014) observed the influence of the average depth on the distribution of this same population of franciscanas. Amaral et al. (2018) also identified depth as limiting the distribution of the species, without detecting slope effects. However, this study used wide spatial scales to assess the topographic slope of studied environments, which may have weakened the power of analysis of this variable. In two gulfs in southern Australia, the habitat use of the bottlenose dolphin (*Tursiops truncatus*) is also associated with a flat bottom topography (Bilgmann et al. 2019).

The heterogeneous presence of franciscanas within Babitonga Bay was found to be associated with sand in the bottom sediment. The species occurs mainly in coastal regions, outside bays and estuaries, where sandy bottoms predominate, and although the Babitonga’ population is exclusively resident in an estuarine environment (Cremer & Simões-Lopes 2008), it may still maintain preferences related to the habitual distribution of the species. The preference of sandy bottom areas by franciscanas has already been noted, especially in the spring, with an increase in the use of muddy areas in winter (Paitach et al. 2017). These findings were based on visual sightings, but they are now corroborated and expanded by the present study. When we look at the foraging areas at dawn and night, there was an increase in the use of muddy bottom areas, demonstrating that these areas are also important for the population in the spring. A very similar result was observed for the harbour porpoise in the Moray Firth, Scotland, where only sandy banks were identified as important foraging areas without including time variables (Brookes et al. 2013), but when the diel cycles were investigated, adjacent muddy areas were also found to be important habitats for them at night (Williamson et al. 2017).

Studies on the habitat use of franciscanas throughout its distribution are rare, partly explained by the difficulty of studying this species in the wild. Based on bycatch data, Danilewicz et al. (2009) observed that the distribution of franciscanas in Rio Grande do Sul reaches predominantly up to 30 meters in depth. That study, however, did not investigate whether water depth is an important factor related to the distribution of the species. More recently, Amaral et al. (2018) analyzed the influence of environmental variables to predict the spatial niche of franciscanas on a wide scale, verifying that depth and salinity can be limiting. Using aerial surveys of distribution over a wide area in southeastern and southern Brazil, Sucunza et al. (2019) observed 54 groups of franciscanas in waters with an average depth of 7.15 m. Although focused on a typical estuarine population, the novel habitat use investigation presented here allow insights into important environmental features to the species in general.

The influence of environmental cycles on cetaceans is mainly a consequence of variations in the availability of their prey (Hastie et al. 2004). Predators must be able to take advantage of these temporal changes in the aquatic environment to optimize feeding success (Lin et al. 2013). However, the distribution dynamics between predators and prey are bidirectional—both sides in this relationship affect each other— so predators seek to optimize prey capture and prey correspondingly to reduce risk of predation (Trites 2009, Becker & Suthers 2014). Thus, the trade-off between foraging success and predator avoidance is decisive in the habitat use of a species (Trites 2009). The franciscanas have no frequent predators in Babitonga Bay, such as large sharks and orcas (Cremer 2015, Gerhardinger et al. 2020). Therefore, the availability of prey is the main factor affecting its distribution. Franciscana is considered a generalist and opportunistic species, preying on the most abundant small fish species in the environment (Cremer et al. 2012, Paitach 2015). However, considering the bidirectionality of the predators-prey relationship mentioned above, it is expected that competing predators will affect each other, a subject that will be discussed below in the specific session on the sympatry between the franciscana and the Guiana dolphin.

Despite contributing to improving the model’s AIC, it is not clear how many of the factor variables are related to the presence of franciscanas. Many levels were not precisely estimated, as indicate by large p-values (Supp. material). The modelling approach adopted here was adequate to provide insights into the environmental variables related to the occurrence of franciscanas within Babitonga Bay. However, model fit was not perfect, although optimal with the selected variables, and therefore, this ecological investigation could greatly benefit from further modelling exploration, such as: inclusion of additional variables (e.g., prey availability), exploring more complex interactions between variables, modelling habitat use for specific periods (e.g., additional seasons), exploring models that accommodate more complex autoregressive structures, etc.

### 4.3. Sympatry with the Guiana dolphin

The intensity of presence of Guiana dolphins was identified in the models as the main variable related to habitat use of franciscanas in Babitonga Bay. Cremer (2007) observed a high overlap in the spatial niche of these populations, but with no competition for interference between them (sensu Bearzi 2005), which has been reaffirmed over the years (Cremer et al. 2018). Analysis of stomach content point to a high degree of prey sharing between the species (Cremer et al. 2012, Paitach 2015). It is interesting to note that although both species have wider amplitudes of the trophic niche in the cold months, when the prey availability is lower (Cremer 2007), there is a decrease in the trophic overlap between them, attenuating the effects of competition (Paitach 2015). This may be the reason why our models showed some overlap between the two species, with franciscana apparently indifferent to the presence of Guiana dolphins up to an extreme extent (Fig. 3).

Different ecological processes may be involved in the niche partition between ecologically similar species living in direct sympatry, such as differences in behavior patterns and diet, differences in habitat use and temporal segregation in the use of resources (Parra et al. 2006, Nichol et al. 2013, Méndez-Fernandez et al. 2013). Considering the high overlap of the trophic and spatial niches, and the absence of agonistic interactions between franciscanas and Guiana dolphins in Babitonga Bay (Cremer et al. 2018), we suggest that the main factors that make possible the coexistence of these two species are fine-scale differences in the habitat use with temporal segregation in the foraging behavior. A fine-scale study of Guiana dolphin’s habitat use and other analytical approaches that integrate different spheres of the realized niche of both species, would assist in elucidating that question.

### 4.4. Spatio-temporal patterns of occurrence and foraging

The distribution of the franciscanas was predominant in the central region of the bay, with greater dispersal in winter than in spring, with virtually no detections in the connection channel with the open sea in either season. This corroborates conclusions from previous studies derived from visual observations (Cremer & Simões-Lopes 2008, Cremer et al. 2018). The Guiana dolphins also have larger areas of distribution in seasons with less prey availability elsewhere (Wedekin et al. 2010) and in Babitonga Bay (Cremer et al. 2011). However, we observed a much more acute use of the center-south portion of the bay in relation to what was observed in previous studies. In fact, franciscana preys are known to concentrate in the region of the bay (Cremer 2007, Paitach 2015). In the present study, the central-southern portion of the distribution area was most frequented at night and at dawn, and mainly for foraging purposes. The innermost muddy banks in the western part of the estuary are also used for foraging, especially on spring afternoons. Since foraging is expected to intensify when/where individuals can maximize their food intake (Pirotta et al. 2014), cyclic of use of such areas can be related with the distribution of the Guiana dolphin. Not surprisingly, the central-southern portion of the bay is considered the core area of Guiana dolphin distribution (Cremer et al. 2011, Cremer et al. 2018).

The present study is the first to analyze the distribution of franciscanas throughout the day and to preliminarily identify the main foraging areas in Babitonga Bay, on seasonal and diel scales. Multiscale approaches have been shown to be very useful in studies of distribution of highly mobile species that explore dynamic habitats (González-García et al. 2018), such as the characteristics of the environment and species dealt with here. In particular, the association of foraging with specific environmental characteristics must be considered in the management of anthropic disorders (New et al. 2013, Pirotta et al. 2014). In the present work, the distribution analyzes were descriptive and did not aim to relate the foraging behavior with environmental characteristics, however such an approach would be desirable in future studies.

### 4.5. Implications for management and conservation

Some anthropogenic activities in Babitonga Bay constitute direct or indirect threats to the survival of this population of franciscanas, such as the overfishing, water pollution, intense vessel traffic and port building and maintenance activities (Cremer 2007, Paitach et al. 2019). Above all, the cumulative and potentially synergetic effects of the different anthropogenic impacts on coastal environments put the dolphins under strong pressure and are often neglected by environmental authorities (Cremer 2007, Azevedo et al. 2017, Herbst et al. 2020). The establishment and operation of big ports represent a major threat to marine biodiversity, causing acute disturbances and a chronic decrease in environmental quality (Domit et al. 2009). Underwater blasting work, periodic dredging of the seabed and intensification of sea traffic result in suspension of sediments and thereby increase the bioavailability of contaminants, oil blades on the surface, increased underwater noise and the risk of collision between cetaceans and vessels, among other impacts that disrupt the natural communities, reduce the availability of prey and compromise the entire health of the ecosystem (Domit et al. 2009, Jefferson et al. 2009, Herbst et al. 2020). It is known that franciscanas avoid areas with known higher levels of underwater noise in Babitonga, which are close to the existing ports (Holz 2014). It has also been observed that after activities requiring the use of dredges, pile drivers and other heavy machinery, the Guiana dolphins abandoned the São Francisco do Sul port inlet for years (Cremer et al. 2018).

Several new ports are planned in Babitonga Bay, of which at least three in the areas identified as critical habitats for the franciscanas. In light of the results presented here, some key aspects must be considered in environmental impact studies, such as: 1) the importance of franciscana foraging areas as critical habitats for their survival; 2) the impacts caused to the population of Guiana dolphins can also result in fundamental consequences for the franciscanas, considering the competitor-predator-prey relationship; 3) the exclusion of artisanal fishing areas, due to the delimitation of the vessels’ maneuvering areas in ports, will cause an intensification of fishing in the center-north portion of the bay, increasing the bycatch risk of franciscanas; and 4) the cumulative and potentially synergistic impacts caused by the new ports added to the ports already operating in the territory.

In Babitonga, dredging for the extraction sand from the bottom is constant throughout the year (Herbst et al. 2020), and the uncontrolled removal of this substrate can also be an indirect threat to the franciscanas, as indicated by the association between the species’ habitat use and this type of substrate found in our study. The operation of dredgers also generates substantial noise, which can be impactful for franciscanas (Holz 2014). The licensing of new sand extraction areas needs to take this potential negative impact into account and adopt the necessary mitigation measures, such as avoiding critical franciscana habitats.

The franciscana bycatch in the artisanal fisheries, although not so frequent in Babitonga Bay, still represents a severe threat considering that the removal of any individual from this small population can be critical to its sustainability (Pinheiro & Cremer 2003, Cremer et al. 2018). Distribution and foraging maps presented here can guide a participatory construction and implementation of exclusion zones in periods of most intense use. Unfortunately, there is no efficient mechanism for fisheries management in the territory, making it difficult to implement strategies to prevent accidental captures, such as fishing exclusion zones or the use of acoustic deterrent devices on nets (FAO 2021).

In recent years, many marine protected areas (MPA) have been designated with the aim of managing human activities for the protection of marine mammals (Hoyt 2012). Dynamic approaches with flexible spatial and temporal limits of protection areas have been recommended for mobile species such as dolphins (Castro et al. 2014, Santos et al. 2017, Hazen et al. 2018, Tardin et al. 2020). However, there are many difficulties for the creation or effective implementation and maintenance of MPA’s in Brazil, such as lack of staff and funding, deficient or absent interinstitutional governance, excessive bureaucracy, and lack of political incentives for any significant change (Gerhardinger et al. 2011). The proposal to create an MPA in Babitonga Bay has been underway in the national environmental agency (i.e., Chico Mendes Institute for Biodiversity Conservation – ICMBio, Brazilian Ministry of the Environment’s) for over ten years (Herbst et al. 2020).

### 4.6. Final considerations

PAM with C-PODs has provided to be a useful method to get important information for management of low density and threatened cetacean populations worldwide, such as the vaquita (*Phocoena sinus*) (Jaramillo-Legorreta et al. 2016), the Maui dolphin (*Cephalorhynchus hectori maui*) (Rayment et al. 2011), the Baltic harbour porpoise (Carlén et al. 2018), and now the franciscanas of Babitonga Bay. Unfortunately for vaquitas that information came too late, and the species may be on the brink of extinction (D’Agrosa et al. 2000). If we don’t act while the franciscana population in Babitonga Bay still is genetically viable, we can see an early extinction of this singular and critically endangered population. This study provides new insights into their habitat use and distribution, identifying the critical habitat for its conservation (sensu Cañadas et al. 2005). The challenge ahead is to identify effective ways to integrate the information on the ecological needs of the franciscana into relevant public policies for the human activities management.

## Acknowledgments

We are very grateful to all the members and collaborators of the Toninhas Project/Univille for their field assistance, especially Ana Bárbara Broni, Gabriel Teixeira, Júlio César dos Santos and Tiago Ramos de Andrade. We are also grateful for the help of Dr. Len Thomas during the sample planning phase of the research. We are grateful to Fundo de Apoio à Pesquisa – FAP/UNIVILLE, Yaqu Pacha Foundation and Petrobras SA for funding this study. To Swedish Agency for Marine and Water Management – SwAM, for lending some of the C-PODs used in this study. To Chelonia Ltd. for donations and all technical support. RLP thanks to Coordenação de Aperfeiçoamento de Pessoal de Nível Superior – CAPES for the PhD research grant. MJC thanks to CNPq for a research productivity scholarship (10477/2017-4).

