## Supplementary material for "Critically endangered franciscana dolphins in an estuarine area: fine-scale habitat use and distribution from acoustic monitoring in Babitonga Bay, southern Brazil": Supp. material

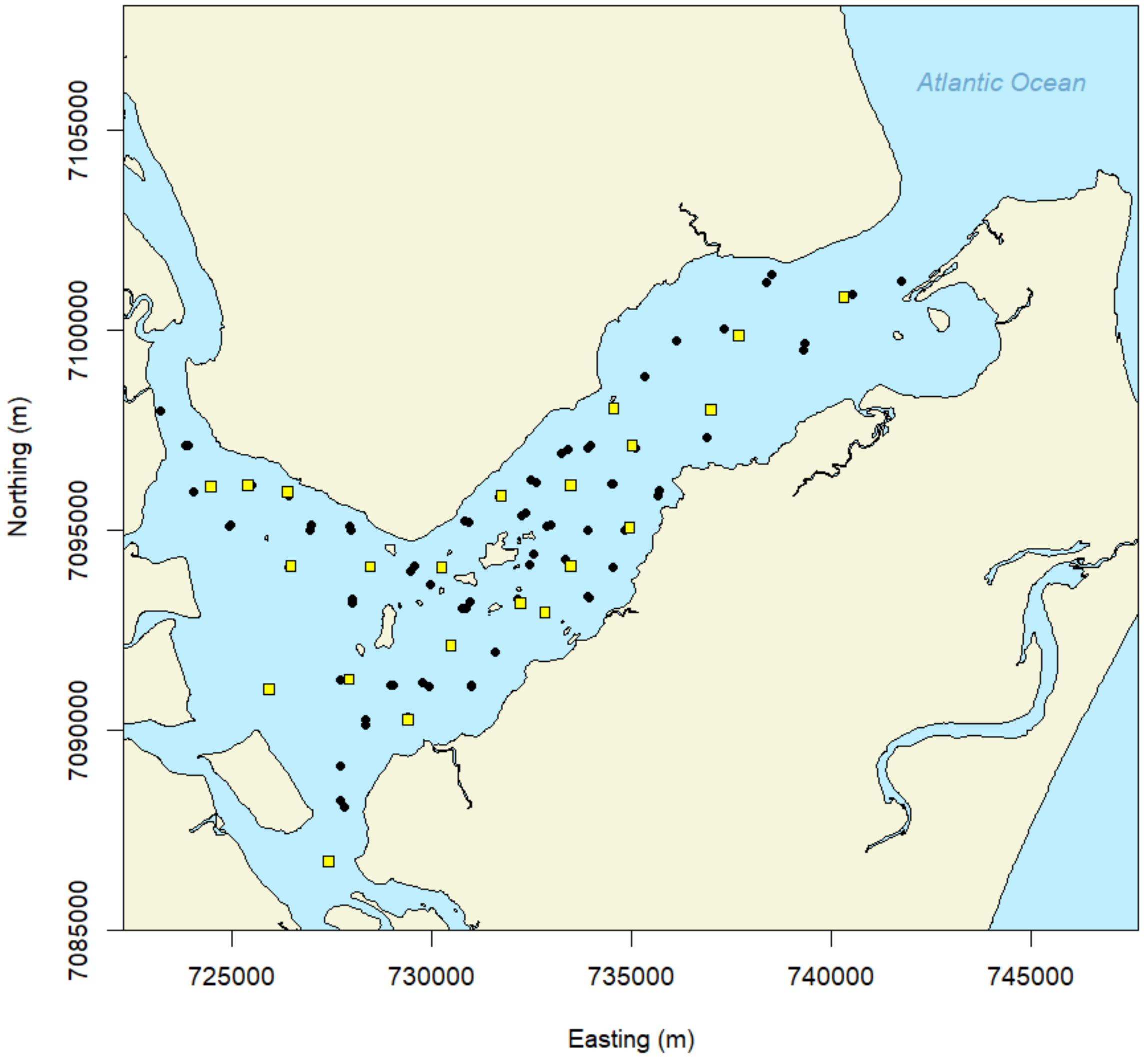


Figure S1. Points acoustically sampled in Babitonga Bay are shown as black dots. Sampling points with less than 100 sampling units (i.e., data points) are overlaid as yellow squares.

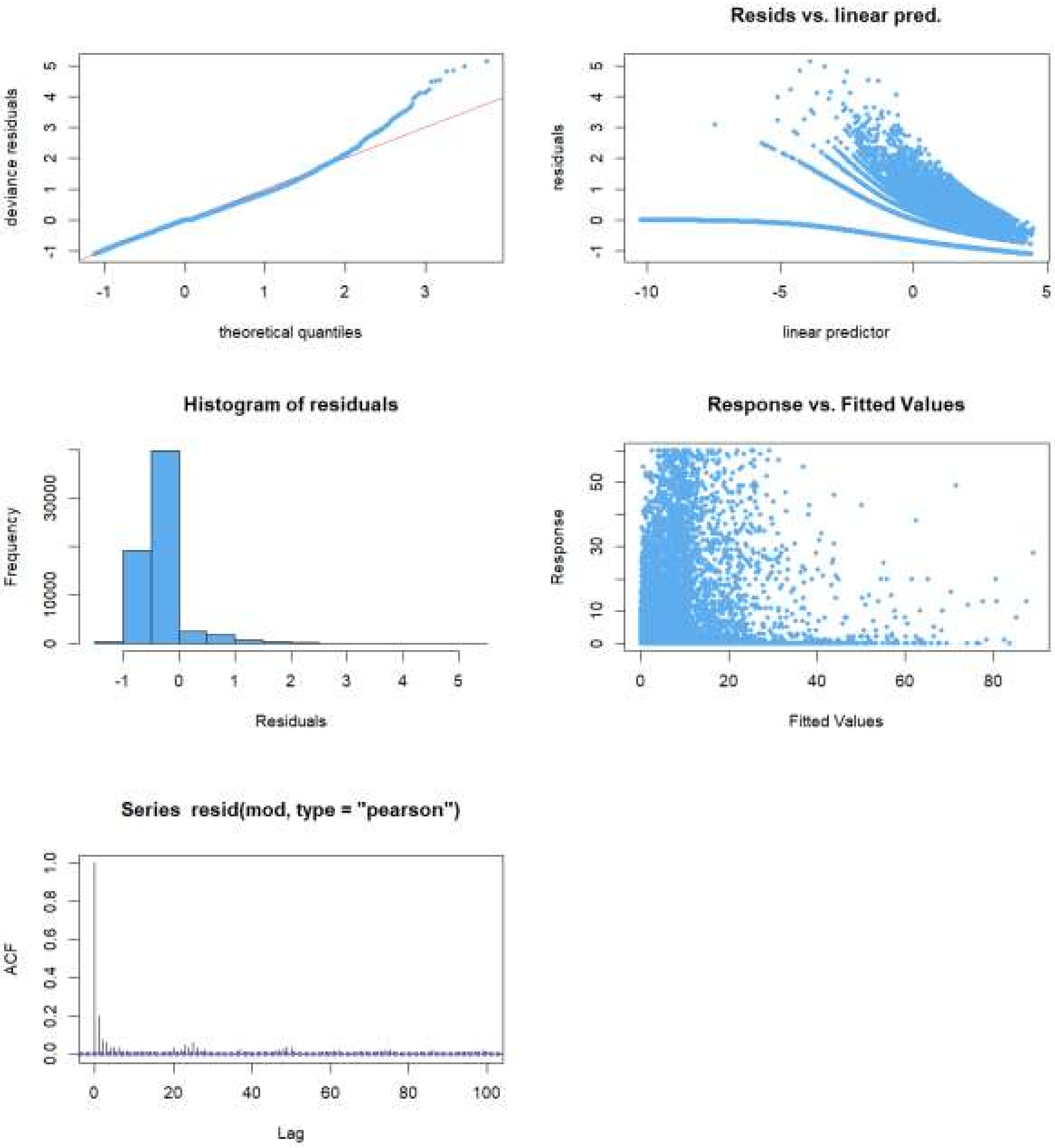


Figure S2. Diagnostic plots for the final model of habitat use of franciscana dolphins in Babitonga Bay.

Table S1. Coefficients for factor variables included in the final model of habitat use of franciscana dolphins in Babitonga Bay.

| **Parametric coefficients** | **Estimate** | **p-value (t-distribution)** |
|---|---|---|
| (Intercept) | -3.253 | < 0.001 |
| Month07 | 0.428 | 0.012 |
| Month08 | 0.685 | < 0.001 |
| Month09 | 0.591 | 0.002 |
| Month10 | 0.119 | 0.516 |
| Month11 | -0.006 | 0.974 |
| Month12 | -0.404 | 0.123 |
| Sed. – mud + sand | 1.830 | < 0.001 |
| Sed. – sand | 0.679 | 0.124 |
| Sed. – sand + mud | 1.776 | < 0.001 |
| Tide.type.syzygy | 0.103 | 0.060 |
| Tide.state.flood | -0.007 | 0.880 |
| Tide.state.high | 0.021 | 0.673 |
| Tide.state.low | -0.073 | 0.147 |

Table S2. Results for smooth functions included in the final model of habitat use of franciscana dolphins in Babitonga Bay. (edf = effective degrees of freedom)

| **Smooth terms** | **edf** | **p-value (F-statistic)** |
|---|---|---|
| s(Point) | 50.843 | < 0.001 |
| s(UTMY,UTMX) - Hour.day 1 | 15.399 | 0.002 |
| s(UTMY,UTMX) - Hour.day 2 | 14.826 | 0.015 |
| s(UTMY,UTMX) - Hour.day 3 | 9.548 | 0.035 |

| | | |
|---|---|---|
| s(UTMY,UTMX) - Hour.day 4 | 18.351 | 0.002 |
| s(UTMY,UTMX) - Hour.day 5 | 8.325 | 0.053 |
| s(UTMY,UTMX) - Hour.day 6 | 14.545 | < 0.001 |
| s(UTMY,UTMX) - Hour.day 7 | 13.102 | 0.008 |
| s(UTMY,UTMX) - Hour.day 8 | 2.000 | 0.060 |
| s(UTMY,UTMX) - Hour.day 9 | 2.001 | 0.085 |
| s(UTMY,UTMX) - Hour.day 10 | 2.001 | 0.011 |
| s(UTMY,UTMX) - Hour.day 11 | 17.015 | < 0.001 |
| s(UTMY,UTMX) - Hour.day 12 | 2.000 | 0.010 |
| s(UTMY,UTMX) - Hour.day 13 | 2.000 | 0.065 |
| s(UTMY,UTMX) - Hour.day 14 | 16.314 | 0.003 |
| s(UTMY,UTMX) - Hour.day 15 | 2.002 | 0.068 |
| s(UTMY,UTMX) - Hour.day 16 | 17.340 | < 0.001 |
| s(UTMY,UTMX) - Hour.day 17 | 2.002 | 0.050 |
| s(UTMY,UTMX) - Hour.day 18 | 2.001 | 0.107 |
| s(UTMY,UTMX) - Hour.day 19 | 16.523 | 0.002 |
| s(UTMY,UTMX) - Hour.day 20 | 12.259 | 0.006 |
| s(UTMY,UTMX) - Hour.day 21 | 10.998 | 0.003 |
| s(UTMY,UTMX) - Hour.day 22 | 9.488 | 0.073 |
| s(UTMY,UTMX) - Hour.day 23 | 11.014 | 0.003 |
| s(UTMY,UTMX) - Hour.day 24 | 11.824 | < 0.001 |
| s(Hour.day) | 3.281 | < 0.001 |
| s(Sg.DPM) | 5.861 | < 0.001 |
| s(Slope.max) | 3.695 | < 0.001 |
